# Site-specific integration of transposon via engineered piggyBac transposase

**DOI:** 10.64898/2026.01.20.700476

**Authors:** Naiqing Xu, Lei Han, Xiaohan Hu, Yuan Fang, Limeng Wu, Xi Wang, Haoyang Tu, Xiaogang Deng, Weili Cong, Kailiang Sun, Yi Jin, Xiaoyang Wu

**Affiliations:** GeCell Therapeutics, LLC, Shanghai, China; Ben May Department for Cancer Research, The University of Chicago, Chicago, IL 60637, USA

## Abstract

The precise integration of large DNA fragments into the human genome holds significant therapeutic potential. Here, we demonstrate that combining engineered piggyBac (PB) transposase with CRISPR/Cas9 enables targeted integration of PB transposons into specified genomic loci. Our engineered PB transposase (PBase) retains high excision activity while substantially reducing endogenous integration activity. In the developed Cas9-PBase fusion system, PBase excises the transposon to generate linear DNA fragments, while Cas9 introduces site-specific double-strand breaks (DSBs), facilitating insertion of the excised fragment at the target locus. The optimized tool achieves 6.1–7.3 kb transposon integration at multiple genomic sites with 10–15% efficiency, demonstrating 60–80% targeted integration specificity. As a proof of concept, we inserted a 7.1 kb transposon encoding three genes into the β2M locus of human induced pluripotent stem cells (iPSCs), conferring protection against allogeneic natural killer (NK) cell-mediated cytotoxicity in derived iNK cells. These results establish Cas9-PBase as a precise and programmable platform for large DNA sequence insertion with potential clinical applications.

## Introduction

The past decade has witnessed remarkable advancements in genome editing technologies, particularly through the development of CRISPR-Cas systems and their derivative tools including base editors (BEs) and prime editors (PEs)[1–4]. These revolutionary tools have enabled unprecedented precision in genomic modifications, transforming research across biological sciences, agriculture, and biomedicine. However, current editors are limited to modifying short sequences, and CRISPR-mediated homology-directed repair (HDR) remains inefficient for inserting large DNA fragments. Despite ongoing tool development and optimization, the precise integration of large DNA sequences into specific mammalian genomic loci continues to present substantial technical hurdles[5].

Human genetic diseases frequently arise from diverse mutational events including single-nucleotide variants, large deletions/insertions, inversions, and translocations. For many such disorders, particularly monogenic diseases like cystic fibrosis (caused by hundreds of mutations distributed across the 250 kb CFTR locus)[6], a gene supplementation strategy involving stable genomic integration of functional transgenes offers distinct advantages over mutation-specific correction approaches. Current integration platforms fall into three main categories: viral vectors, transposon systems, and CRISPR-Cas-mediated homologous recombination[7–9]. While viral and transposon systems demonstrate efficient gene transfer capabilities, their semi-random integration patterns raise significant genotoxicity concerns[10–12]. Conversely, CRISPR-Cas systems enable precise targeting but show limited efficiency for large fragment integration via HDR[13].

The piggyBac (PB) transposon system represents a promising alternative, employing a “cut-and-paste” mechanism that efficiently integrates large DNA fragments flanked by inverted terminal repeats (ITRs) into TTAA sites without leaving footprint sequences[14, 15]. This unique combination of features provides an ideal foundation for developing next-generation integration tools. We hypothesized that engineering a fusion protein combining PB transposase (PBase) with a sequence-specific nuclease could leverage PBase’s efficient large-fragment excision/integration capabilities while achieving precise genomic targeting[16–18]. Critical to this approach is developing PBase variants that maintain robust transposon excision activity while minimizing off-target integration at endogenous TTAA sites[16–18]. Recent studies have explored such Exc^+^Int^-^ mutants[19, 20], suggesting this strategy’s potential for achieving high-efficiency, site-specific integration of large DNA fragments.

A rational approach to minimize PBase-mediated off-target integration at TTAA sites involves structure-guided mutagenesis of DNA-interacting residues to reduce target binding affinity. The previously characterized Exc^+^Int^-^ mutant (PB^R372A_K375A_D450N^) demonstrated this principle by maintaining robust transposon excision while exhibiting attenuated TTAA integration[21]. In this study, the investigators engineered a fusion protein by combining the Exc^+^Int^-^ mutant with a zinc-finger protein (ZFP) domain designed to target specific DNA-binding sequences; however, no substantial integration bias was detected at the genomic loci corresponding to the ZFP recognition sites. Subsequent development of the FiCAT system through Cas9 fusion achieved sgRNA-directed precise integration[19, 20], though with suboptimal efficiency and incomplete off-target profiling.

To overcome these limitations, we propose engineering next-generation PBase variants with enhanced transposon excision activity to increase substrate availability for targeted integration and further suppressed TTAA integration propensity to minimize off-target events. This dual-optimization strategy should simultaneously improve both the efficiency and specificity of programmable DNA insertion.

To develop PBase variants with optimized excision activity and minimized TTAA integration, we performed structural analysis of the PBase-DNA complex during transposition. Previous crystallographic studies identified six critical residues (Y312, R315, L324, N347, K375, R376) within the catalytic and insertion domains that mediate contacts with the DNA backbone flanking TTAA sites[22]. These predominantly polar, positively charged amino acids facilitate electrostatic interactions with the negatively charged phosphate groups of target DNA.

Based on these structural insights, we hypothesized that strategic mutagenesis of these residues to either non-polar alanine or negatively charged glutamic acid would disrupt favorable electrostatic interactions with target DNA, while preserve the enzyme’s ability to recognize and excise transposon DNA. This rational design approach was predicted to specifically attenuate TTAA-dependent integration activity while maintaining or enhancing transposon excision capability.

Through systematic single-mutation analysis of the six candidate residues, we identified N347, K375, and R376 as critical determinants of integration efficiency, with glutamic acid substitutions at these positions dramatically reducing PBase activity. Building on these findings, we engineered a series of double and triple mutants that exhibited either enhanced excision capability and more stringent suppression of TTAA integration compared to the reference Exc^+^Int^-^ mutant (PB^R372A_K375A_D450N^).

Fusion of these optimized PBase variants to *Streptococcus pyogenes* Cas9 enabled precise, sgRNA-directed transposon integration. Through iterative optimization of the Cas9-PBase system, we achieved targeted integration efficiencies of 6-15% for 6.1-7.3 kb transposons, with on-target specificity reaching 60-80%. As a functional demonstration, we successfully employed this system to simultaneously disrupt the β2M locus and integrate a 7.1 kb triple-gene construct in human induced pluripotent stem cells (iPSCs) via single-step editing. These results establish our engineered Cas9-PBase platform as an efficient and precise tool for therapeutic genome engineering applications, combining high-capacity DNA delivery (≥7 kb) and reduced off-target integration.

## Results

### Development of PBase mutants with high transposon excision activity or/and low TTAA-dependent integration activity

Structural analysis of the PBase-DNA complex[22] identified six critical amino acid residues (Y312, R315, L324, N347, K375, and R376) within the catalytic and insertion domains that mediate interactions with the DNA backbone flanking TTAA recognition sequences (Figure 1B). To systematically evaluate the functional contribution of each residue to transposition activity, we generated a comprehensive series of single-point mutants by substituting each position with either non-polar alanine (A) or negatively charged glutamic acid (E) residues.

**Figure 1.**
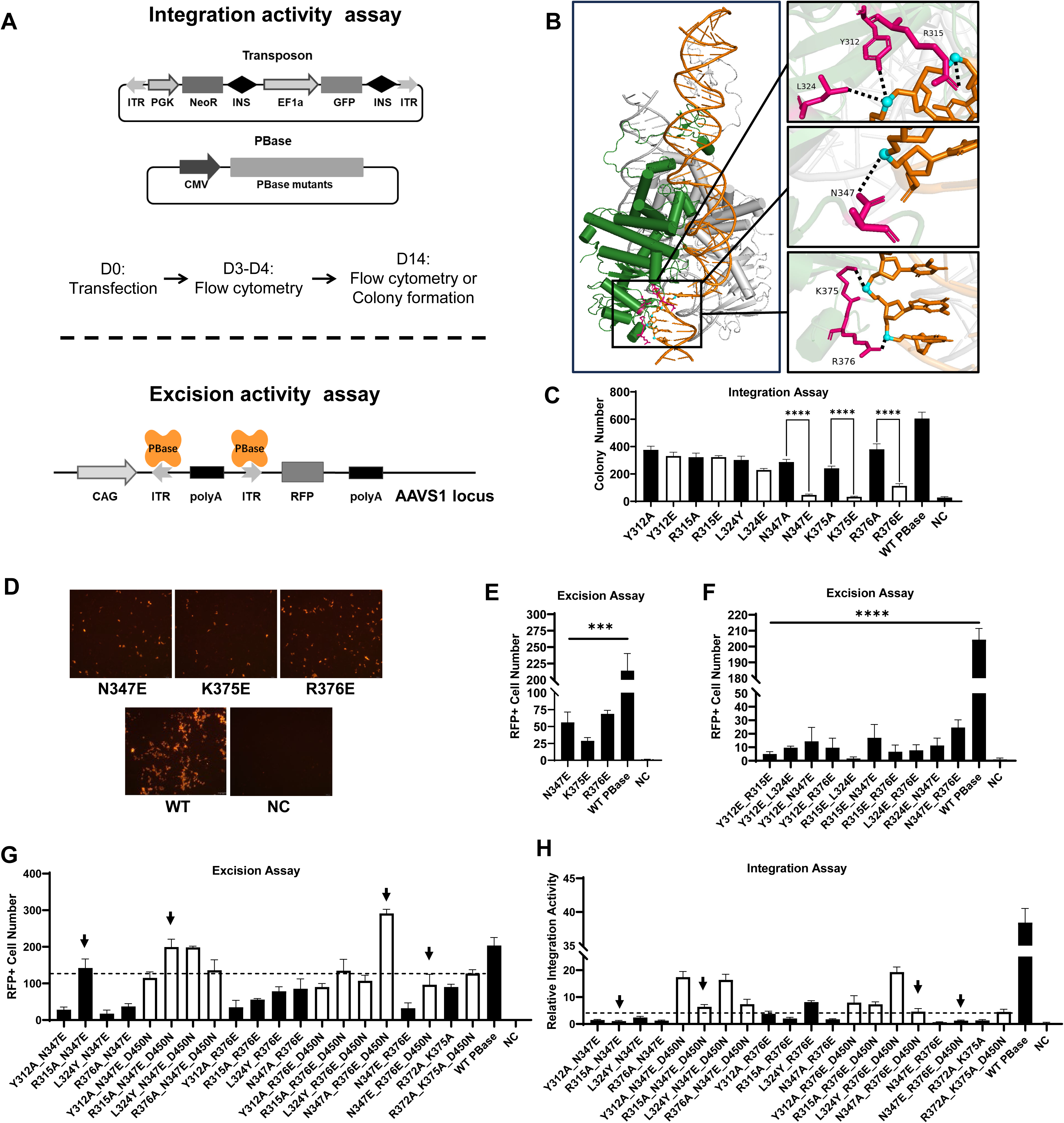
Integration and excision activity of PBase mutants in human cells. (A) Schematic illustration of the integration assay system (upper panel) and the excision assay system (lower panel). (B) Detailed interactions of PBase amino acids (Y312, R315, L324, N347, K375 and R376) with phosphates in the target DNA. (C) Integration activity of single mutants measured by the colony formation assay. G418 resistant colonies were counted and represent the integration activity of indicated PBase mutant. The negative control (NC) is background integration without PBase and the positive control is integration by a wild-type (hyperactive) PBase. (D) Measuring the excision activity of PBase mutants by restoring RFP expression. The number of RFP+ cells represents the excision activity of indicated PBase mutant. The negative control (NC) is background fluorescence without PBase. Representative image of n = 3. (E) The RFP positive cell counts from transfection of single PBase mutants, and the wild-type PBase respectively. (F) The RFP positive cell counts from transfection of double PBase mutants containing two glutamic acid substitutions. (G) The RFP positive cell counts from transfection of indicated double and triple PBase mutants. Arrows indicate selected mutant candidates. (H) The integration activity of indicated double and triple mutants. Integration activity is represented by stable expression rate of GFP cargo (D14) normalized to transfection efficiency(D3). Arrows indicate selected mutant candidates. All data was analyzed and presented as Mean ± SD of three biological replications, *p≤0.05, **p≤0.01, ***p≤0.001, ****p≤0.0001.

We employed a colony formation assay in HeLa cells to quantitatively assess the integration activity of each PBase variant. Cells were co-transfected with two plasmids: a *piggyBac* transposon construct encoding GFP and neomycin resistance markers (Figure 1A, upper panel), and expression vectors for either wild-type PBase (a hyperactive variant designated as wild-type for reference) or individual mutant forms. Quantitative analysis revealed that alanine substitutions at Y312, R315, N347, K375, and R376 positions or tyrosine substitutions at L324 resulted in partially reduced integration activity, which maintained 40-70% of wild-type as measured by colony formation. Similar activity profiles were observed for the Y312E, R315E, and L324E mutants. In striking contrast, glutamic acid substitutions at N347, K375, and R376 positions dramatically impaired transposition efficiency. The PB^K375E^ mutant exhibited the most severe phenotype, with colony formation rates approaching background levels observed in transposase-negative controls. The N347E and R376E mutants showed slightly higher residual activity (7.6% and 18.6% of wild-type, respectively) than K375E, but still markedly reduced compared to both wild-type and corresponding alanine variants (Figure 1C). These findings establish N347, K375, and R376 as critical determinants of PBase integration activity, with glutamic acid substitutions at these positions preferentially disrupting transposition function.

To precisely evaluate the excision capabilities of our lead candidates (N347E, K375E, and R376E), we established a sensitive reporter system in HEK293T cells. Using homology-directed repair (HDR), we integrated a reporter construct into the AAVS1 safe harbor locus. This system features a CAG promoter-driven Red Fluorescent Protein (RFP) gene whose expression is suppressed by an intervening *piggyBac* transposon cassette containing a polyadenylation signal (Figure 1A, lower panel). Successful transposon excision by PBase restores RFP expression, providing a quantitative measure of excision activity (Figure 1D).

Validation experiments confirmed the system’s reliability, with wild-type PBase transfection producing robust RFP expression. Among the glutamic acid mutants, K375E demonstrated severely impaired excision capacity (13% of wild-type activity), while N347E and R376E retained moderate activity (26-32% of wild-type; Figure 1D and 1E). Given the requirement for efficient excision in our application, we excluded K375E from further development despite its strong integration suppression. N347E and R376E emerged as potential candidate mutants, although further reductions of their integration activity was required.

To achieve optimal Exc^+^Int^-^ properties, we investigated combinatorial mutations. Initial attempts with double glutamic acid substitutions yielded variants with minimal excision activity, with PB^N347E_R376E^ showing relatively higher RFP expression, at approximately 12% of wild-type PBase (Figure 1F). These results indicate that most double E PBase mutants are not suitable for further study, except for PB^N347E_R376E^. Parallel characterization of double alanine mutants (PB^Y312A_R315A^, PB^Y312A_L324Y^, PB^Y312A_R376A^, PB^R315A_L324Y^, PB^R315A_R376A^, and PB^L324Y_R376A^) revealed robust colony formation in integration assays (data not shown), indicating insufficient suppression of off-target activity. These results guided our subsequent engineering of more sophisticated triple mutants.

Next, we introduced a second alanine substitution into the N347E or R376E single mutants. We hypothesized that these E_A or A_E double mutants might exhibit higher excision activity than E_E mutants while maintaining reduced integration activity compared to the original N347E or R376E single mutants. To test this, we engineered eight E_A or A_E double mutants derived from N347E and R376E. Furthermore, given that the D450N mutation has been previously reported to substantially enhance excision activity in Exc^+^Int^-^ mutants[21], we incorporated this mutation into our double mutants to generate corresponding triple mutants. We then systematically assessed the excision and integration activities of these double and triple mutants.

Comparative analysis revealed that, relative to the PB^R372A_K375A_D450N^ benchmark mutant, four novel mutants demonstrated either higher or comparable excision activity alongside similar or reduced integration activity. These included PB^R315A_N347E^, PB^R315A_N347E_D450N^, PB^N347A_R376E_D450N^ and PB^N347E_R376E_D450N^. Notably, the PB^N347A_R376E_D450N^ mutant displayed approximately 2.3-fold higher excision activity than the benchmark mutant while maintaining comparable integration efficiency, suggesting its potential utility for enhanced transposon excision in site-specific integration applications. In contrast, other mutants, such as PB^N347E_R376E_D450N^, exhibited 75% of the reference mutant ’s excision activity and only 28% of its integration activity, implying a reduced propensity for off-target integrations at TTAA sites (Figure 1G, 1H). Collectively, these findings indicate that the four newly identified Exc^+^Int^-^ mutants represent promising candidates for the development of improved site-specific integration tools.

### Site-specific integration of Cas9-PBase mutants in a split-GFP system

To engineer a site-specific integration tool, we genetically fused candidate PBase mutants to Streptococcus pyogenes Cas9 via a 3×G4S flexible linker and evaluated the resulting fusion enzymes’ ability to direct transposon integration at predetermined genomic loci. To quantitatively assess the efficiency and targeting specificity of Cas9-PBase-mediated integration, we implemented a split-GFP reporter system in HeLa cells, adapted from established methodologies[19, 23]. This system enables emGFP expression exclusively upon successful site-specific integration events. The reporter cell line was generated by stably integrating a promoterless C-terminal GFP fragment (C-GFP) containing the target sequence into the genome. The 6.1 kb donor transposon construct contained a CAG promoter-driven N-terminal GFP fragment (N-GFP) and a constitutively expressed BFP-P2A-PuroR selection cassette. Functional GFP reconstitution only occurs when the transposon integrates in correct orientation at the target site, thereby serving as a precise readout for successful integration events (Figure 2A).

**Figure 2.**
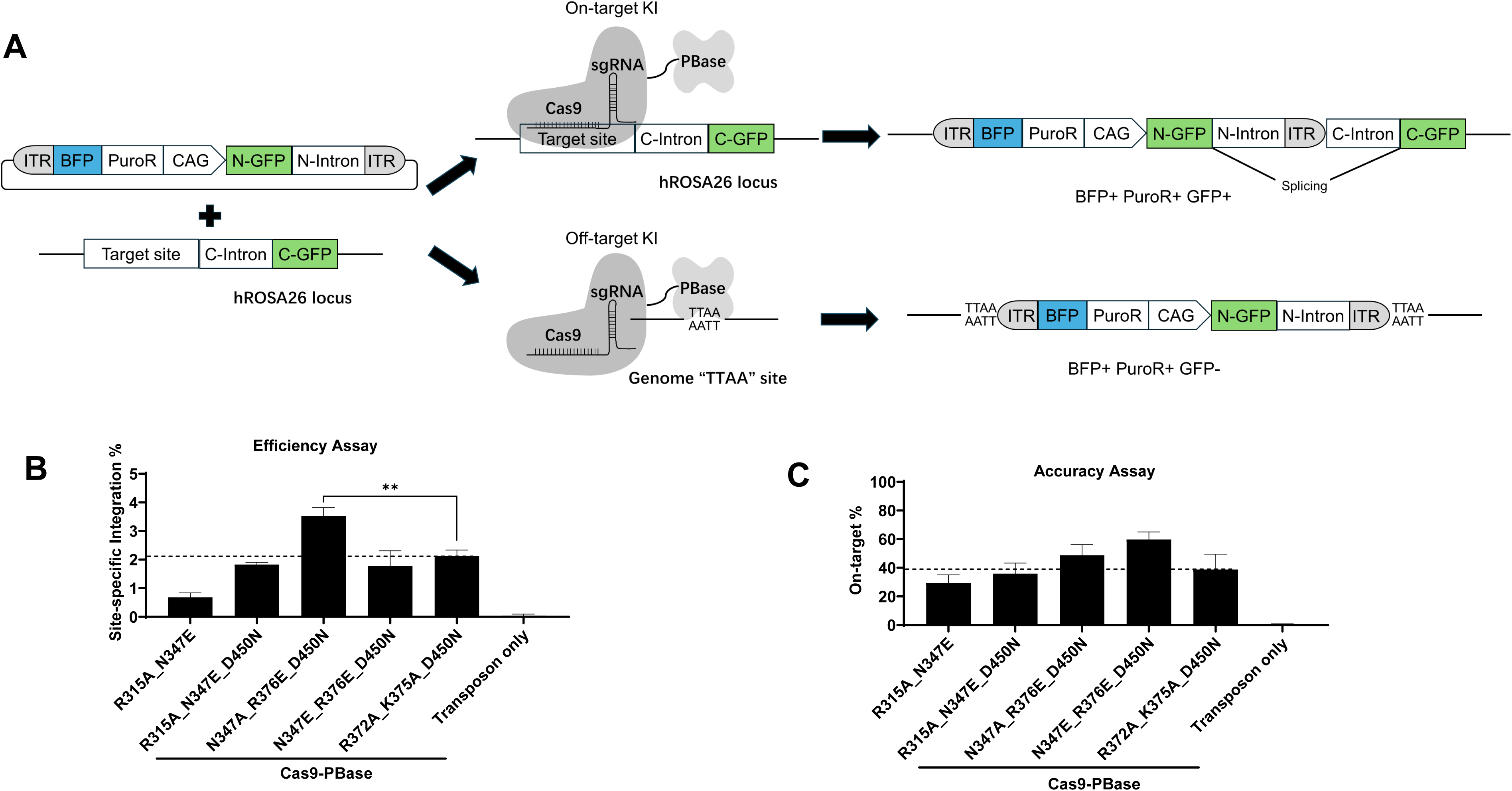
Site-specific integration mediated by Cas9-PBase fusion protein. (A) Illustration of the split-GFP reporter system to test site-specific integration by Cas9-PBase. With this reporter, the transposon donor expressing BFP, PuroR and N-terminal GFP is directed to the target site upstream of C-GFP by Cas9-PBase. BFP positive cell counts represent the transfection rate (Day 4) or total integrated rate (Day14). GFP positive cell counts represent forward on-target integration. (B) Site-specific integration efficiency mediated by selected Cas9-PBase mutants. The ratio of 2 × GFP/BFP(D4) represents the site-specific integration efficiency. Cas9-PBase R372A_K375A_D450N is used as the benchmark. (C) The on-target ratio of selected Cas9-PBase mutants. Stable integrated cells were enriched with puromycin and analyzed by flow cytometry at day 14. The on-target percentage is represented as 2×GFP/BFP(D14). All data was analyzed and presented as Mean ± SD of three biological replications, *p≤0.05, **p≤0.01, ***p≤0.001, ****p≤0.0001.

For targeting experiments, we constructed plasmids expressing Cas9-PBase fusions and sgRNAs targeting the hROSA26 locus (target sequence: GTCGAGTCGCTTCTCGATTA) upstream of the C-GFP sequence. The N-GFP donor plasmid and Cas9-PBase-sgRNA constructs were co-transfected into the C-GFP reporter cell line. At 4 days post-transfection, we quantified GFP+ (successful integration) and BFP+ (total transfected) populations by flow cytometry. Since transposons can integrate in either orientation, we calculated site-specific integration efficiency as 2×GFP+/BFP+ to account for both possible configurations.

Comparative analysis revealed that the Cas9-PB^N347A_R376E_D450N^ fusion exhibited ∼60% higher integration efficiency than the benchmark Cas9-PB^R372A_K375A_D450N^, while Cas9-PB^N347E_R376E_D450N^ and Cas9-PB^R315A_N347E_D450N^ showed comparable efficiency to the reference (Figure 2B). Following puromycin selection (day 14), we further evaluated integration precision by calculating the on-target ratio (2×GFP+/BFP+). The Cas9-PB^N347A_R376E_D450N^ mutant (48.7% on-target) demonstrated similar precision to Cas9-PB^R372A_K375A_D450N^ (38.8%), while Cas9-PB^N347E_R376E_D450N^ (59.7%) showed superior targeting accuracy (Figure 2C).

These findings correlate well with our earlier characterization of TTAA integration activity (Figure 1H), where PB^N347E_R376E_D450N^ exhibited reduced off-target integration compared to PB^R372A_K375A_D450N^. Collectively, these results demonstrate that Cas9-PB^N347A_R376E_D450N^ represents an improved variant that combines enhanced integration efficiency with maintained targeting precision relative to the benchmark PB^R372A_K375A_D450N^ mutant.

### Cas9-PBase mediated integration at endogenous genome loci

To assess the performance of our engineered Cas9-PBase system in unmodified genomic contexts, we conducted a comprehensive evaluation at three well-characterized safe harbor loci: hROSA26, GSH1, and GSH2[24]. We designed twelve distinct Cas9-PBase-sgRNA constructs targeting these sites, incorporating four variants: the reference Cas9-PB^R372A_K375A_D450N^ and three optimized mutants (N347A_R376E_D450N, N347E_R376E_D450N, and R315A_N347E_D450N). Each locus was targeted using validated sgRNA sequences (hROSA26: GTCGAGTCGCTTCTCGATTA; GSH1: TTAGTCATGCATGCATGAAG; GSH2: CATCAGACTTGATAGATAGCATG).

In HeLa cells co-transfected with the 7.3 kb GFP-NeomycinR transposon donor and respective Cas9-PBase-sgRNA plasmids (Figure 3A), we observed distinct integration profiles across variants. Quantitative flow cytometry analysis revealed that Cas9-PB^N347A_R376E_D450N^ achieved superior integration efficiency, demonstrating 2.04-fold and 1.89-fold higher GFP expression at GSH1 and GSH2 loci, respectively, compared to the benchmark. While Cas9-PB^R315A_N347E_D450N^ showed comparable or slightly reduced efficiency, Cas9-PB^N347E_R376E_D450N^ exhibited moderately lower integration rates at GSH loci while maintaining high specificity (Figure 3B). These findings align with our previous results from the split-GFP reporter system.

**Figure 3.**
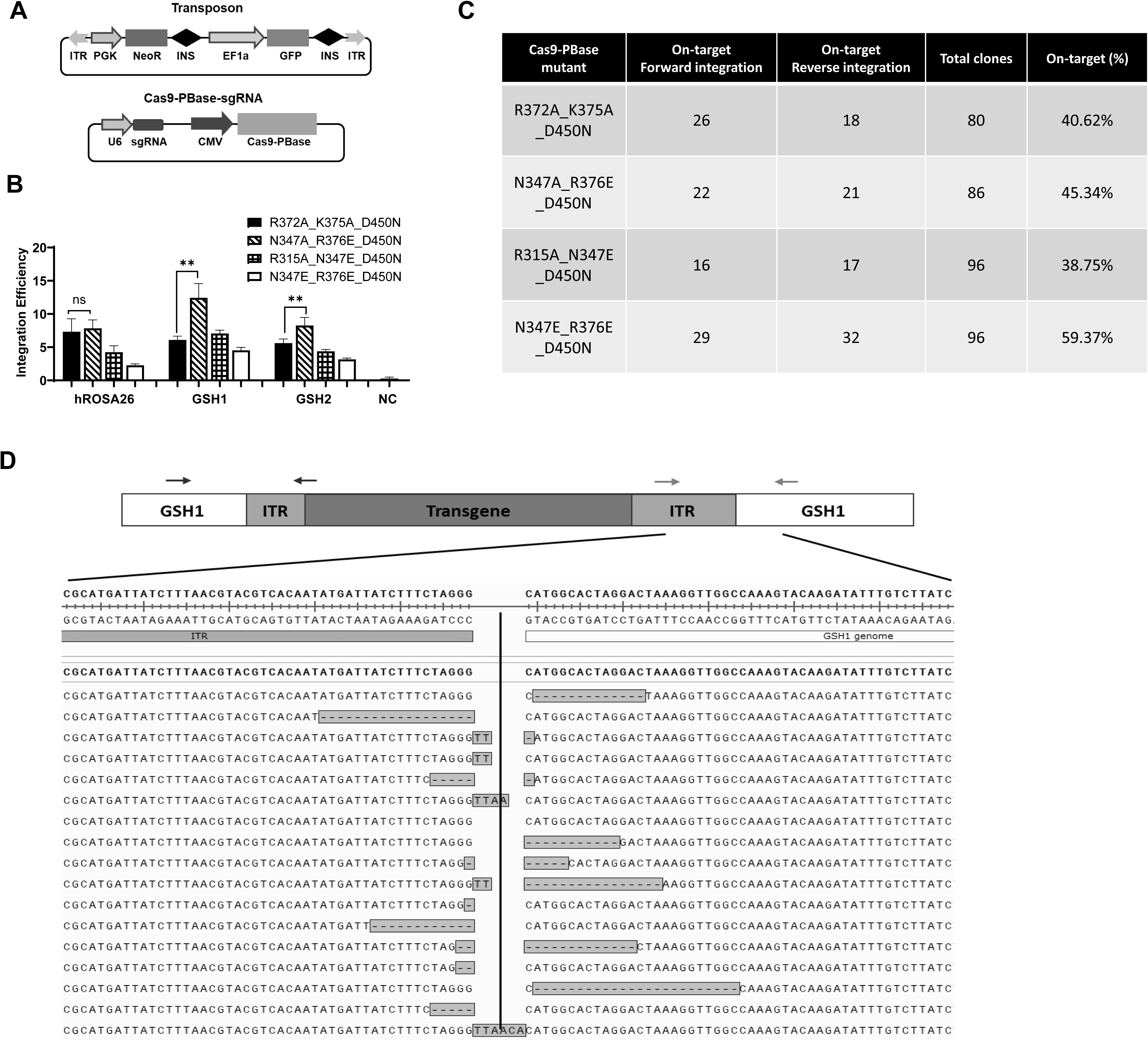
Characterization of Cas9-PBase mutants for endogenous genomic loci. (A) Two plasmids system to test the site-specific integration of piggyBac transposon at endogenous genomic loci. The transposon plasmid expresses NeoR & GFP. The enzyme plasmid expresses Cas9-PBase and transcribes sgRNA for target site. (B) Total integration efficiency of Cas9-PBase at endogenous genomic loci. Integration efficiency is represented by GFP(D14)/GFP(D3) in HeLa cells. The negative control (NC) is background integration without Cas9-PBase enzyme. (C) On-target clones at GSH1 locus determined by junction-PCR. Single cell clones were digested and lysed for junction PCR with primer pairs annealing to genomic loci and transposon donor respectively. On-target clones were determined with PCR positive bands. (D) Characteristics of junction sequences between ITR and genomic DNA. Representative sequencing reads of on-target clones are aligned to the reference sequence. Small insertions or deletions were found in the junction, suggesting the transposon sequence is integrated in HITI manner. Arrows indicate two primer pairs for junction PCR. All data was analyzed and presented as Mean ± SD of three biological replications, *p≤0.05, **p≤0.01, ***p≤0.001.

Detailed molecular characterization of GSH1 integrations through single-cell cloning and junction-PCR analysis of 86 clones generated with Cas9-PB^N347A_R376E_D450N^ revealed balanced orientation distribution (22 forward, 21 reverse, and 4 biallelic integrations). Comparative analysis of on-target rates demonstrated that Cas9-PB^N347E_R376E_D450N^ achieved the highest specificity (59.37%), followed by Cas9-PB^N347A_R376E_D450N^ (45.34%) and the benchmark (40.62%). Cas9-PB^R315A_N347E_D450N^ showed the lowest on-target rate (38.75%) (Figure 3C). Analysis of 358 total clones confirmed equivalent frequencies of forward (93) and reverse (88) integrations, indicating unbiased orientation preference during the integration process.

Sanger sequencing of over 100 junction-PCR products revealed characteristic small indels at integration sites, consistent with the homologous-independent target integration (HITI) mechanism[25]. Representative sequence alignments of forward integration clones showed precise insertion at the DSB site with minimal flanking modifications (Figure 3D). These results collectively demonstrate that Cas9-PB^N347A_R376E_D450N^ combines enhanced integration efficiency with maintained precision, while operating through a predictable HITI mechanism that shows no orientation bias. The system’s performance at endogenous loci confirms its potential for reliable genomic engineering applications.

### Optimization to improve the efficiency and accuracy of the Cas9-PBase system

Building upon our initial success with the Cas9-PB^N347A_R376E_D450N^ variant (V1), we pursued additional engineering strategies to further enhance the system’s performance. Based on structural insights revealing key residues (N286 and T557) involved in ITR interactions[22] (Figure 4A), we hypothesized that strengthening PBase’s binding affinity to transposon ends could improve delivery of linear DNA fragments to target sites. Introduction of positively charged arginine substitutions at these positions yielded significant improvements: the single-mutation variant Cas9-PB^N347A_R376E_D450N_T557R^(V2) demonstrated 3.05-fold higher integration efficiency than the benchmark, while the double-mutant Cas9-PB^N286R_N347A_R376E_D450N_T557R^(V3) achieved a remarkable 4.55-fold enhancement (Figure 4B-C).

**Figure 4.**
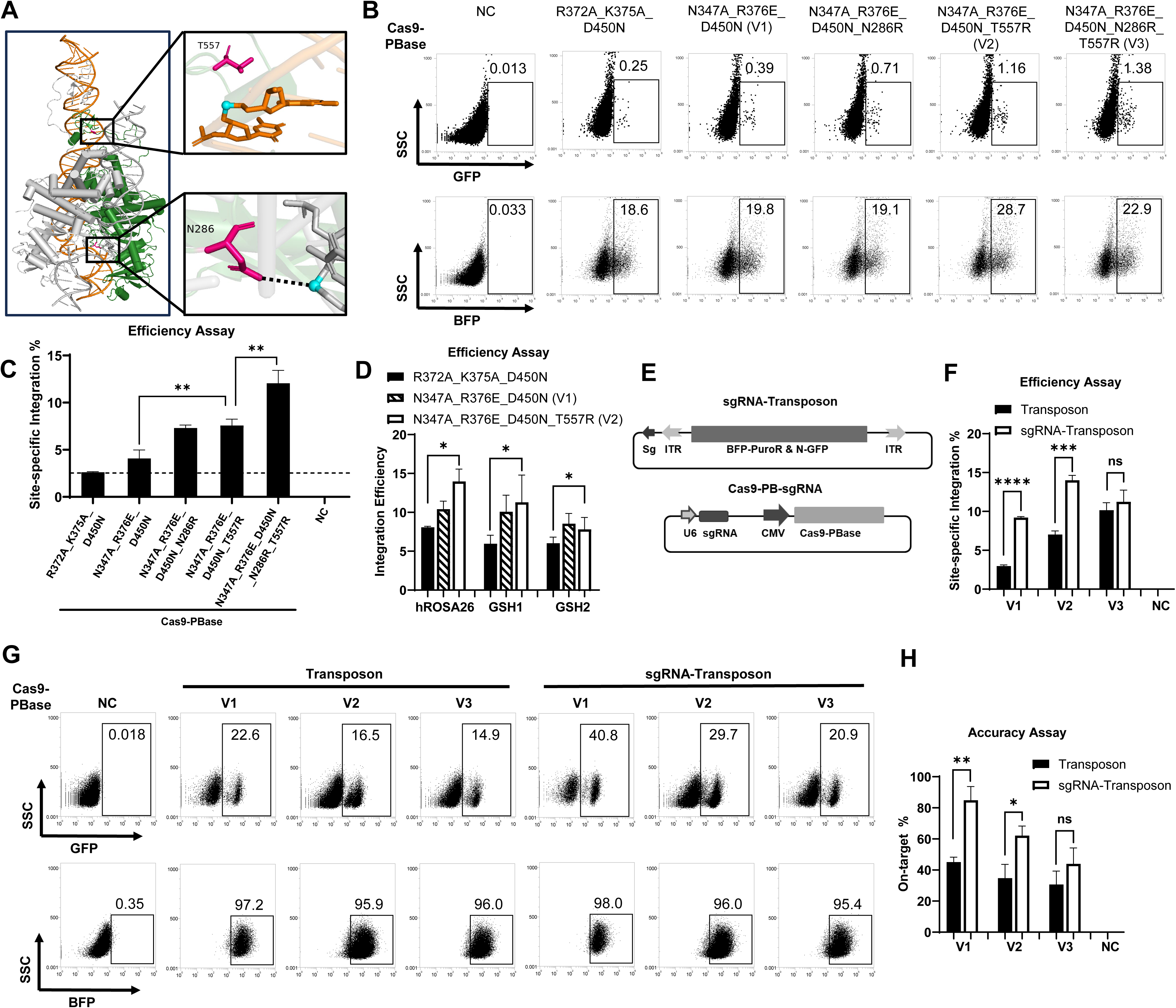
Optimization to improve efficiency and accuracy of the Cas9-PBase system. (A) Detailed interactions between N286/T557 of PBase with phosphates in the ITR DNA. (B) Flow cytometry analysis of further engineered Cas9-PBase in the split-GFP system on day 4 post transfection. Cas9-PBase N347A_R376E_D450N is termed as v1, Cas9-PBase N347A_R376E_D450N_T557R is v2, and Cas9-PBase N347A_R376E_D450N_N286R_T557R is v3. The negative control (NC) is background fluorescence of reporter cells. Representative image of n = 3. (C) Efficiency of site-specific integration by further engineered Cas9-PBase. The ratio of 2×GFP/BFP (D4) represents the site-specific integration efficiency. (D) Total integration efficiency of Cas9-PBase at endogenous genomic loci. Integration efficiency is represented by GFP(D14)/GFP(D3) in HEK293T cells. Background integration without Cas9-PBase has been deducted. (E) Schematic illustration of the sgRNA-Transposon plasmid with sgRNA target sequences located upstream of the 5’ ITR. (F) Efficiency of site-specific integration of sgRNA-Transposon by Cas9-PBase versions in the split-GFP system. The ratio of 2×GFP/BFP (D4) represents the site-specific integration efficiency. (G) Flow cytometry analysis of puromycin-enriched cells on day 14 post transfection. Stable integrated cells in Figure 4E were selected with puromycin. The negative control (NC) is background fluorescence without transfection. Representative image of n = 3. (H) On-target ratio of the sgRNA-Transposon and Transposon by Cas9-PBase variants. The ratio of 2×GFP/BFP (D14) in Figure 4F represents the on-target percentage. All data was analyzed and presented as Mean ± SD of three biological replications, *p≤0.05, **p≤0.01, ***p≤0.001, ****p≤0.0001.

Validation in HEK293T cells at endogenous loci (hROSA26, GSH1, GSH2) confirmed these improvements, with V2 showing 12-34% higher efficiency than V1, and both variants substantially outperforming the original benchmark (Figure 4D). Notably, V2 achieved 10-15% integration efficiency for 7.3 kb transposons at safe harbor sites, representing a significant advancement in large DNA fragment delivery.

We implemented an additional optimization strategy by modifying the transposon donor architecture to improve both integration efficiency and targeting accuracy. Specifically, we incorporated an sgRNA target sequence adjacent to the 5’ TTAA-inverted terminal repeat (ITR) of the transposon donor, creating a novel “sgRNA-Transposon” construct (Figure 4E). This design builds upon previously established approaches that demonstrated enhanced homologous recombination efficiency through similar modifications to DNA templates[26, 27]. The modified sgRNA-Transposon exhibits two key functional advantages over the conventional design: First, it becomes susceptible to dual recognition-capable of being cleaved and bound by either Cas9 or PBase, whereas the unmodified transposon can only be processed by PBase. This dual recognition mechanism potentially enhances integration efficiency by increasing the probability of donor DNA engagement. Second, Cas9’s physical occupancy near the 5’ ITR may sterically hinder PBase-mediated integration at random TTAA sites, thereby improving targeting specificity.

We systematically compared the performance of the sgRNA-Transposon versus the original construct using our split-GFP reporter system. Quantitative analysis revealed that the sgRNA-Transposon mediated significantly higher GFP expression rates when paired with both V1 and V2 enzyme variants compared to the conventional transposon (Figure 4F), demonstrating markedly improved site-specific integration efficiency. Notably, the V2 variant achieved a 14% integration efficiency (normalized to transfection efficiency) with the 6.1 kb sgRNA-Transposon (Figure 4F).

Following puromycin selection of stably integrated cells, we performed comprehensive precision assessments via flow cytometry. The sgRNA-Transposon showed substantial improvements in targeting accuracy across all tested variants: V1 increased the GFP+/BFP+ cell ratio from 22% to 40%, V2 enhanced it from 16.5% to 29.7%, and V3 improved it from 14.9% to 20.9% (Figure 4G). Most impressively, the calculated on-target integration ratio (2×GFP+/BFP+) reached 60-80% for both V1 and V2 when using the sgRNA-Transposon (Figure 4H).

Collectively, these optimizations yielded a Cas9-PBase system with approximately 4-fold greater site-specific integration efficiency compared to our original benchmark (Cas9-PB^R372A_K375A_D450N^), while simultaneously maintaining or improving targeting precision. These improvements establish the sgRNA-Transposon as a superior donor design for precise genome engineering applications.

### Precise large cargo integration by Cas9-PBase in hiPSCs for stable expression

To evaluate the clinical potential of our optimized Cas9-PBase system, we applied the V2 variant (Cas9-PB^N347A_R376E_D450N_T557R^) to human induced pluripotent stem cells (hiPSCs), focusing on developing immune-compatible iNK cell therapeutics. The challenge of allogeneic rejection in iNK cell therapy requires simultaneous knockout of MHC-I (via β2M disruption) and knock in of protective genes (HLA-E and CD47) to evade host immune surveillance[28, 29]. Given the narrow editing window in hiPSCs due to their limited expansion capacity before differentiation, we designed a 7.1 kb transposon containing three therapeutic elements to achieve these modifications in a single step.

Using a β2M-targeting sgRNA (GGCCACGGAGAGACATCT), we co-delivered the transposon donor and Cas9-PBase V2 constructs to hiPSCs (Figure 5A). HLA-E-positive cells were enriched via antibody-coated microbeads, yielding 12 correctly targeted clones out of 43 screened (28% efficiency). Genotypic characterization revealed six clones with biallelic modification (KI/KO) and six with monoallelic modification (KI/wt). Whole-genome sequencing of selected KI/KO clone #29 (carrying an 8-bp β2M exon 1 deletion) confirmed precise single-copy integration at the target locus without detectable off-target events.

**Figure 5.**
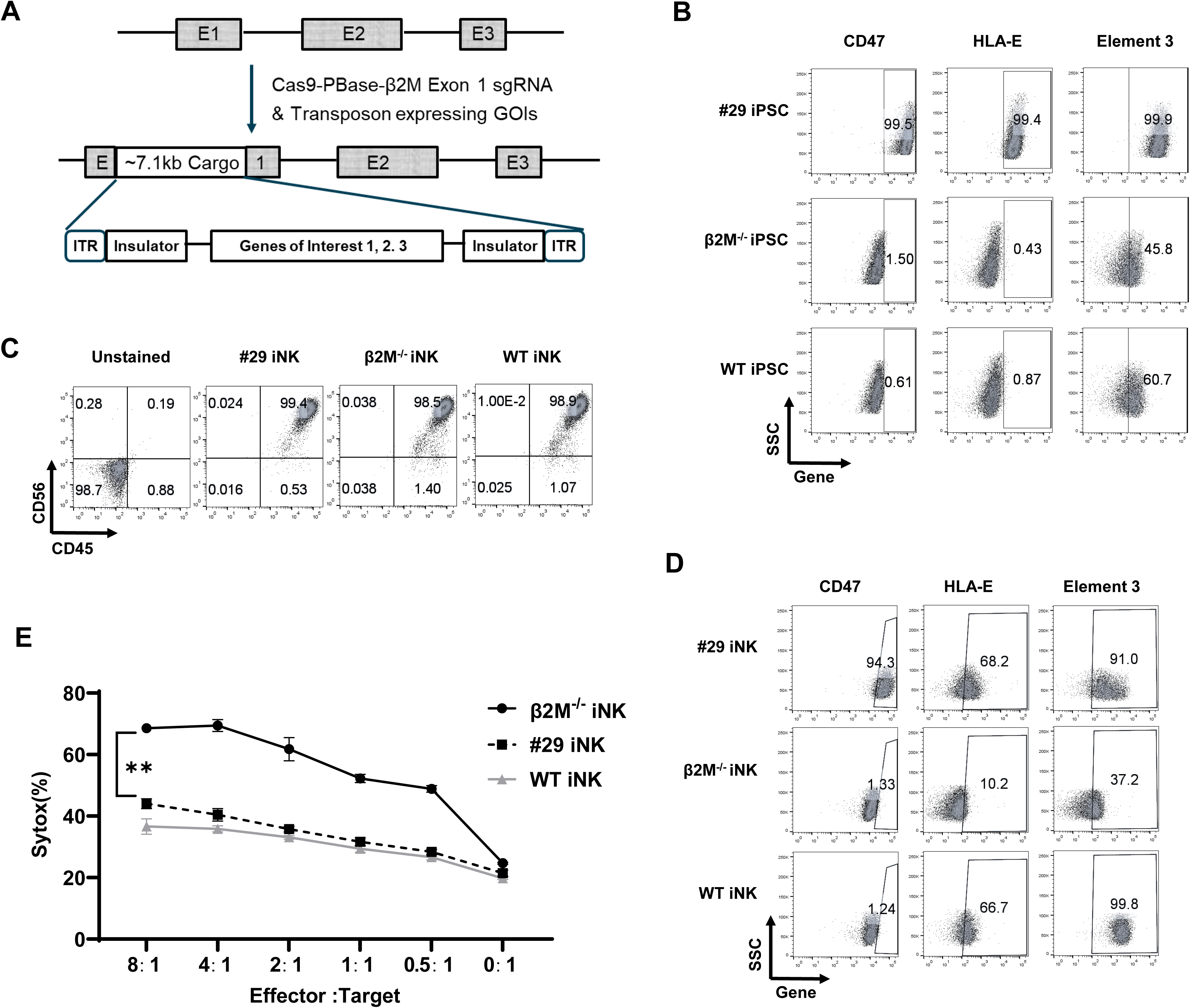
Large cargo integrated precisely by Cas9-PBase in hiPSC. (A) Strategy of iPSC editing by Cas9-PBase. The sgRNA was designed to target exon 1 of β2M gene, allowing knock-out of β2M and knock-in of the 7.1 kb cargo within a single editing step. (B) Transgenes expression quantification by flow cytometry analysis of edited iPS cells. The edited #29 iPSC clone, the β2M^-/-^ control clone and the wild-type iPSCs were stained with antibodies to detect expression of CD47, HLA-E and element 3. (C) Characterization of iNK cells. Differentiated iNK cells at day 28 of differentiation stage 2 were stained with CD45 and CD56 antibodies, followed by flow cytometry analysis. (D) Transgenes expression quantification by flow cytometry analysis of iNK cells. Differentiated iNK cells at day 28 of differentiation stage 2 were stained with antibodies to detect expression of CD47, HLA-E and element 3. (E) Cytotoxicity by allogeneic PBNK to iNK cells. Prelabeled iNK cells were co-cultivated with allogeneic PBNK cells at multiple E:T ratios for 6 hours. The cytotoxicity of PBNK to iNK cells was measured with SYTOX staining and flow cytometry analysis. Representative image of n = 3. All data was analyzed with Mean ± SD of three biological replications, *p≤0.05, **p≤0.01, ***p≤0.001.

Flow cytometric analysis demonstrated robust transgene expression in edited hiPSCs, with clone #29 showing significantly higher CD47 and HLA-E mean fluorescence intensity (MFI) compared to both wild-type and β2M-/- controls (>99% positivity for all transgenes; Figure 5B). Following a two-stage differentiation protocol (Zhu and Kaufman, 2019), the edited hiPSCs successfully generated iNK cells with >98% purity (CD56+CD45+) comparable to unmodified controls (Figure 5C). Importantly, transgene expression persisted throughout differentiation, with iNK cells from clone #29 maintaining elevated CD47 and HLA-E levels versus β2M-/- controls (Figure 5D).

Functional validation through co-culture assays with allogeneic peripheral blood NK cells (PBNK) demonstrated the therapeutic efficacy of our approach. While β2M-/- iNK cells showed increased susceptibility to PBNK-mediated killing compared to wild-type, clone #29 iNK cells exhibited significant protection (Figure 5E), confirming that coordinated β2M knockout with HLA-E and CD47 expression effectively shields iNK cells from host immune responses. In summary, these experiments show that the Cas9-PBase tool can achieve site-specific integration of large DNA fragments in hiPSCs, enabling multi-gene knock-ins and β2M knockout in a single editing step. The edited iPSCs successfully differentiated into high-purity iNK cells with stable expression of the integrated gene elements, functionally reducing PBNK-mediated killing of iNK cells.

## Discussion

Through structure-guided engineering of the PBase-DNA interaction interface, we developed a series of Exc^+^Int^-^ mutants featuring strategically placed non-polar alanine or negatively charged glutamic acid substitutions that significantly reduce DNA binding affinity while maintaining robust excision activity. These engineered variants outperform previously reported Exc^+^Int^-^ mutants in both functionality and precision. When adapted into Cas9-PBase fusion tools, our lead candidate Cas9-PB^N347A_R376E_D450N^(V1) demonstrated 60% greater integration efficiency compared to the benchmark Cas9-PB^R372A_K375A_D450N^. Further optimization through the T557R substitution yielded Cas9-PB^N347A_R376E_D450N_T557R^(V2), which achieved a remarkable 3.5-fold enhancement in integration efficiency. The system’s performance was substantially improved by engineering the transposon donor to include an external sgRNA targeting sequence, enabling 10-15% site-specific integration efficiency for 6.1kb transposons while boosting the on-target ratio from ∼40% to 60-80%. The therapeutic potential of this platform was conclusively demonstrated through efficient single step editing of human iPSCs, simultaneously knocking out β2M while introducing three protective transgenes, ultimately generating iNK cells with enhanced resistance to allogeneic NK cell cytotoxicity.

Structural characterization of the PBase-DNA complex[22] identified six key DNA-interacting residues in PBase, including four highly conserved residues (Y312, R315, N347, K375) and two with lower conservation (L324, R376)[21]. Our systematic analysis demonstrated that N347, K375, and R376 are particularly critical for transposition activity, though previous studies had exclusively focused on K375 modifications. The neutral charge of N347, despite its conservation and proximity to the catalytic D346 residue, along with the variable conservation of positively-charged R376, likely contributed to their prior neglect in mutagenesis studies. Structural analysis revealed these residues engage with negatively-charged DNA phosphates, motivating our innovative strategy of introducing negatively-charged glutamic acid substitutions, an approach previously unreported for PBase-DNA interface engineering. This rational design yielded N347E, K375E, and R376E single mutants showing substantially reduced transposition activity relative to alanine-substituted variants, confirming the functional importance of these residues in DNA binding and transposition.

Building upon our findings with single mutants, we systematically investigated double mutation combinations to optimize the balance between excision and integration activities. While single E-substituted mutants (e.g., N347E and R376E) retained residual integration capacity, we observed that most E_E double mutants demonstrated severely compromised excision activity, falling below the threshold required to generate sufficient linear transposon DNA for efficient site-specific integration. Conversely, A_A double mutants frequently maintained substantial TTAA-dependent integration activity (data not shown), resulting in undesirable off-target events. Through comprehensive characterization, we identified that strategic combinations of A and E substitutions in double mutants achieved our desired profile of robust excision coupled with minimal TTAA-directed integration. Although the current study examined a limited set of double mutants, these proof-of-concept results suggest that expanded screening of combinatorial mutants through systematic library approaches could yield further improvements in system performance.

Analysis of integration junction sequences revealed characteristic small insertions and deletions (indels), consistent with a homologous-independent target integration (HITI) mechanism[25]. We propose the following molecular mechanism for Cas9-PBase-mediated integration: (1) the PBase domain excises the transposon and binds inverted terminal repeats (ITRs), with attenuated integration activity minimizing TTAA-dependent events; (2) simultaneously, the Cas9 moiety generates targeted double-strand breaks (DSBs) guided by sgRNA; (3) the excised transposon integrates at these programmed DSBs during endogenous repair processes (Figure 6). This HITI mechanism offers particular advantages for in vivo applications, as it functions efficiently in non-dividing cells where homologous recombination is largely inactive[25]. These collective findings position Cas9-PBase as a highly promising platform for therapeutic genome editing in post-mitotic tissues.

**Figure 6.**
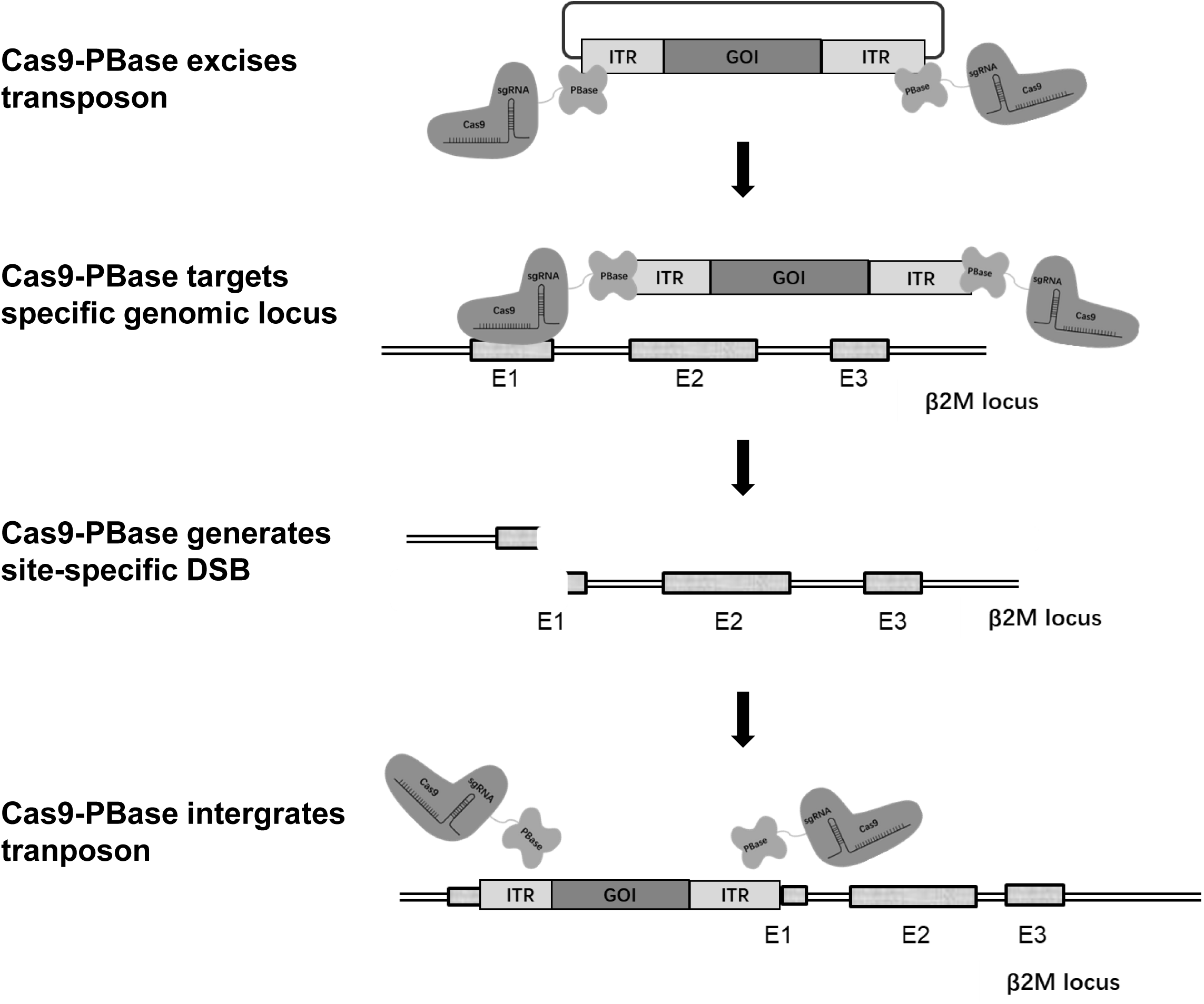
Illustration of Cas9-PBase mediated efficient site-specific integration. PBase in the fusion protein with high excision activity cleaves the transposon and binds to ITR sequences. Since mutated PBase has low integration activity, the transposon payload is not integrated at the “off-target” TTAA sites. Instead, it is directed to the target site by sgRNA, where Cas9 generates the DSB. The payload captured by Cas9-PBase fusion protein to the vicinity of DSB can be integrated in an HITI manner during the repair process.

The development of efficient and precise methods for large DNA fragment integration continues to represent a fundamental challenge in genome engineering. Recent advances have yielded several innovative platforms capable of inserting full-length functional genes, including CAST transposases[30], FiCAT systems[19, 20], PASTE technology[31], PrimeRoot editors[32], PASSIGE platforms[33], dCas9-SSAP fusions[34], zinc-finger recombinases[35], and Bridge RNA-guided recombinases[36]. Within this expanding toolkit, our Cas9-PBase system offers distinct advantages as a streamlined single-enzyme solution that accomplishes transposon integration in a single step while maintaining compatibility with mammalian cell engineering. The platform’s versatility is further underscored by potential optimization through comprehensive PBase mutant screening and transposon architecture engineering. As the field progresses, strategic integration of Cas9-PBase with complementary editing technologies and continued innovation in delivery methods will undoubtedly accelerate therapeutic development for genetic disorders and enhance biopharmaceutical manufacturing capabilities.

## Materials and Methods

### Plasmid Construction

All PBase variants and mutants were cloned into the pcDNA3.1 mammalian expression vector (Invitrogen). The Cas9-PBase fusion constructs were generated using hCas9_D10A (Addgene #41816) as the backbone, with the D10A nickase mutation reverted to wild-type D10 to restore full nuclease activity. A synthetic insert containing a 3×(G4S) flexible linker followed by the PBase coding sequence was ligated to the C-terminus of Cas9 to create the Cas9-PBase expression vector. For sgRNA delivery, synthesized U6-sgRNA cassettes were cloned into MfeI/MluI restriction sites to generate complete Cas9-PBase-sgRNA constructs. Additional plasmids including the GFP-NeomycinR transposon donor, N-GFP transposon, C-GFP acceptor, HLA-C-P2A-CD47-T2A-HLA-E transposon, and RFP reporter plasmids were commercially synthesized (GenScript) and sequence-verified. Complete plasmid sequences and maps are available upon request.

### Cell Culture and Transfection

All cell lines used in this study were maintained under standard conditions with regular mycoplasma testing. HeLa cells (kindly provided by Professor Wu Jian, Fudan University) and HEK293T cells (Chinese National Collection of Authenticated Cell Cultures) were cultured in DMEM (Biosharp) supplemented with 10% fetal bovine serum (Yeasen) and 1% penicillin-streptomycin at 37°C with 5% CO_2_. Selection media contained either 1 μg/mL puromycin (ThermoFisher) or 600 μg/mL G418 (Gibco). Human induced pluripotent stem cells (iPSCs, Shanghai East Hospital) were maintained in TeSR™-AOF medium (STEMCELL Technologies) under feeder-free conditions. Transfections for HeLa and HEK293T cells were performed using Lipofectamine 3000 (Invitrogen) according to the manufacturer’s protocol, while iPSC transfections utilized the Neon™ Transfection System (ThermoFisher) with optimized electroporation parameters.

### PBase Integration Activity Test

Integration activity was assessed using a dual-reporter system in HeLa cells. Cells in 6-well plates were co-transfected with 1.0 μg of GFP-NeomycinR transposon plasmid and 0.7 μg of PBase expression plasmid. GFP expression was quantified by flow cytometry at days 3 and 14 post-transfection, with the D14/D3 GFP+ ratio representing relative integration efficiency. Colony formation assays were performed by plating 5,000 transfected cells (day 10 post-transfection) in 6 cm dishes under G418 selection (600 μg/mL) for 14 days, followed by crystal violet staining.

### PBase Excision Activity Test

To evaluate PBase excision activity, we first generated a stable reporter cell line by precisely integrating an RFP reporter construct into the AAVS1 safe harbor locus of HEK293T cells using homology-directed repair (HDR). The donor DNA template, designed to target the AAVS1 site, contained 5’ and 3’ homology arms flanking the reporter cassette. Following PCR amplification and purification of the donor fragment from genomic DNA, we prepared ribonucleoprotein (RNP) complexes by incubating 150 pmol of Cas9 protein (GenScript, Z03469-500) with 300 pmol of AAVS1-targeting sgRNA (target sequence: CACCCACAGGGGGCCACT) for 10 minutes at room temperature. The RNP complex was then co-electroporated with 1.0 μg of linearized donor DNA into 5×10^6 HEK293T cells using optimized electroporation parameters (1100V, 20ms/pulse, 2 pulses). The integrated reporter construct included a splicing acceptor (SA) site and T2A-Puro cassette driven by the endogenous PPP1R12C promoter. Puromycin selection (1 μg/mL) was applied to isolate single-cell clones, with successful knock-in events verified by Sanger sequencing. This validated reporter cell line subsequently served as the platform for assessing PBase mutant excision activity, where 1 μg of mutant PBase plasmid was transfected using Lipofectamine 3000, followed by quantification of tdTomato-positive cells five days post-transfection.

### Split-GFP Principle and Reporter Cell Line Construction

The reporter system employs mRNA splicing-mediated reconstitution of Emerald GFP (emGFP) expression as a sensitive readout for site-specific integration events. The emGFP coding sequence was strategically divided into two fragments: (1) an N-terminal fragment (N-GFP) incorporated into the transposon donor plasmid, containing both a constitutive promoter and a downstream splice donor site, along with a BFP-P2A-PuroR cassette for monitoring transfection efficiency, quantifying integration events, and selecting stably integrated cells; and (2) a promoterless C-terminal fragment (C-GFP) featuring an upstream splice acceptor site, which was genomically integrated at the ROSA locus to generate the acceptor cell line. Critical to this design is an engineered target sequence upstream of the C-GFP splice acceptor that can be specifically recognized by guide RNAs. Functional emGFP expression occurs exclusively when the N-GFP transposon integrates at the target site adjacent to C-GFP, enabling precise splicing and translation of the full-length reporter.

For reporter cell line establishment, we first introduced the pPGK-Bxb1 attP sequence into the hROSA26 locus (sgRNA target: GTCGAGTCGCTTCTCGATTA) of HeLa cells using Cas9 RNP-mediated homology-directed repair, following similar electroporation and clonal selection protocols as described for the RFP reporter system. Successful knock-in clones were identified by Sanger sequencing and next-generation sequencing (NGS), with preference given to clones exhibiting attP integration at one allele accompanied by indels at the other hROSA26 allele. Subsequent integration of the C-GFP acceptor sequence was achieved by co-transfecting 1.5 μg of the C-GFP donor plasmid with 1 μg of pcDNA3.1-Bxb1 integrase plasmid (derived from Addgene #127519) into the pPGK-attP cell line. Blasticidin selection (10 μg/mL) was applied to isolate correctly integrated clones, which were further validated by Sanger sequencing before establishing a monoclonal acceptor cell line for subsequent experiments.

### Testing Cas9-PBase Integration with the Split-GFP System

For precise measurement of Cas9-PBase integration efficiency, acceptor cells were co-transfected with 1.3 μg Cas9-PBase-sgRNA plasmid (targeting hROSA26) and 1.2 μg N-GFP transposon donor using Lipofectamine 3000. GFP and BFP expression were analyzed by flow cytometry at day 4, with BFP+ cells indicating successful transfection and GFP+ cells indicating site-specific integration. Since transposons can integrate in either orientation, the on-target integration rate was calculated as 2×(GFP+/BFP+) to account for both possibilities. At day 14, puromycin selection (1 μg/mL) was applied to enrich stably integrated cells, followed by flow cytometric analysis to determine the final on-target ratio (2×GFP+/BFP+). This comprehensive approach allowed simultaneous assessment of transfection efficiency, integration efficiency, and targeting specificity throughout the experimental timeline.

## Contributions

Naiqing Xu, Conception and design, Acquisition, Analysis and interpretation of data, Drafting or revising the article; Lei Han, Analysis of data, Revising; Xiaohan Hu, Acquisition and Analysis of data; Yuan Fang, Acquisiton of data; Limeng Wu, Acquisition of data; Xi Wang, Acquisition of data; Haoyang Tu, Acquisition of data; Xiaogang Deng, Acquisition of data; Weili Cong, Revising the article; Kailiang Sun, Conception and design; Yi Jin, Scientific discussion, Revising the article, Supervision, Xiaoyang Wu, Conception and design, Revising the article; Tian Xu, Conception and design, Supervision, Analysis and interpretation of data, Drafting and revising the article.

## Acknowledgement

We thank Professor Tian Xu in School of Life Sciences, Westlake University for giving us useful advice. We thank Professor Wu Jian in School of Basic Medical Sciences, Fudan University for providing HeLa cells.

